# Engineering tunable synthetic fibrotic niches to model acinar-to-ductal metaplasia in 2D pancreatic culture systems

**DOI:** 10.64898/2026.10.15.759386

**Authors:** Shanmukha Subrahmanyam Ayyalasomayajula, Jennifer P. Morton, Catherine C Berry, Matthew J Walker

## Abstract

Acinar-to-ductal metaplasia (ADM) is an early pancreatic cell-state transition associated with injury, inflammation, and progression toward pancreatic intraepithelial neoplasia (PanIN). Although genetic and inflammatory drivers of ADM are well established, the contribution of extracellular-matrix mechanics is difficult to isolate because tissue stiffening is accompanied by changes in matrix composition and soluble signaling. Here, we describe a two-dimensional polyacrylamide (PAA) hydrogel platform that independently varies substrate stiffness and adhesive ligand presentation for murine pancreatic acinar 266-6 cells. Hydrogels approximate soft pancreatic tissue (0.50 ± 0.21 kPa), a transitional mechanical condition (4.48 ± 0.15 kPa), and fibrotic tissue (18.29 ± 0.88 kPa). RGD provides a conventional adhesive cue relevant across healthy-to-fibrotic extracellular-matrix states, whereas incorporation of laminin-derived isoleucine-lysine-valine-alanine-valine (IKVAV) creates a more basement-membrane-like environment consistent with preservation of acinar identity. The platform was also adapted to include gelatin, a collagen-derived protein used here to model a more collagen-rich fibrotic matrix context. Peptide coupling is calibrated using fluorescently labeled RGD, and responses after 3 days are assessed by morphology, alpha-amylase 2 (AMY2), Yes-associated protein (YAP) nuclear-to-cytoplasmic localization, and sex-determining region Y-box transcription factor 9 (SOX9). Increasing stiffness increased nuclear YAP localization and reduced AMY2 expression. IKVAV-containing conditions supported greater attachment on softer matrices and partially preserved acinar-like phenotype on stiff gels despite elevated nuclear YAP. Across all ligand conditions, increasing stiffness reduced roundness and geometric regularity/solidity and increased aspect ratio; on gelatin-modified hydrogels, these phenotypic changes corresponded to increased cell-area-normalized SOX9 fluorescence. Together, these readouts identify a stiffness-dependent shift from an acinar-associated state toward an ADM-associated ductal phenotype, while ligand context modulates how this mechanical response is expressed. This protocol provides a controllable, adaptable system for separating mechanical and biochemical extracellular-matrix inputs during pancreatic phenotypic plasticity.

**SUMMARY:** This protocol describes a tunable two-dimensional polyacrylamide hydrogel platform that independently controls substrate stiffness and extracellular-matrix ligand presentation to model mechanosensitive acinar-to-ductal metaplasia in murine pancreatic acinar cells.

## INTRODUCTION

Pancreatic ductal adenocarcinoma (PDAC) remains one of the most lethal common malignancies. Global estimates for 2022 reported more than 510,000 new pancreatic cancer cases and approximately 467,000 deaths, and five-year survival in the United Kingdom remains below 7% [1,2]. A major reason for this poor outcome is late diagnosis, because early disease is frequently asymptomatic and many patients present only after local invasion or metastatic spread [3]. PDAC develops through a multistep process rather than an abrupt transformation. Mature acinar cells can undergo acinar-to-ductal metaplasia (ADM) following pancreatic injury or inflammation, losing features of digestive-enzyme-producing acinar identity and adopting duct-like characteristics. ADM can be reversible, but persistent inflammatory signaling together with oncogenic events such as Kirsten rat sarcoma viral oncogene homolog (KRAS) mutation can stabilize the transition and support progression through pancreatic intraepithelial neoplasia (PanIN) toward invasive disease [4,5].

The extracellular matrix (ECM) is not only a biochemical scaffold but also a mechanical regulator of cell state. Cells sense substrate stiffness through integrin-based adhesions, the actin cytoskeleton, and mechanotransductive signaling pathways that alter transcriptional programs and cell fate [6-9]. In other tissues, increasing matrix rigidity can reinforce focal adhesion signaling, cytoskeletal tension, and disease-associated phenotypes [8]. The transcriptional co-activators Yes-associated protein (YAP) and transcriptional co-activator with a postsynaptic density protein 95/discs large/zonula occludens-1-binding motif (TAZ) are central components of this response; increased rigidity and cell spreading promote YAP/TAZ nuclear localization, whereas softer environments favor cytoplasmic retention [9]. In the pancreas, YAP/TAZ activity is implicated in loss of acinar identity and acquisition of ductal characteristics during ADM [10]. Chronic pancreatitis, which combines sustained inflammation with progressive fibrotic remodeling, is also a strong risk factor for pancreatic cancer [11]. Importantly, direct mechanical characterization of human pancreatic specimens has shown that both chronic pancreatitis and PDAC tissue are mechanically distinct from histologically normal pancreas, with increased steady-state modulus in both disease states and distinct viscous signatures [12]. This places substantial mechanical remodeling at the pancreatitis stage, a clinically important risk state that can precede neoplastic progression, and supports the possibility that altered mechanics may influence pancreatic cell phenotype before invasive PDAC is established. Hydrogel-based pancreatic models have therefore been highlighted as valuable tools for isolating cell mechanosensing from the complex biochemical changes that accompany disease [13], while broader work on cancer and fibrosis emphasizes that both static stiffness and time-dependent matrix mechanics can regulate pathological cell-state transitions [14]. Recent work from our group further demonstrated that viscoelastic changes between healthy and PanIN-bearing pancreatic tissue can be recreated using two-dimensional polyacrylamide (PAA) hydrogels and that the viscous component of these PanIN-relevant mechanics regulates epithelial-to-mesenchymal transition (EMT)-associated phenotypic changes [15]. Together, these observations support a continuum of mechanically regulated phenotypic plasticity across early pancreatic disease and motivate investigation of whether mechanical remodeling can influence an even earlier transition, ADM.

Mechanical and biochemical ECM changes occur simultaneously in vivo, making them difficult to separate experimentally. Acinar cells are normally associated with basement-membrane-rich environments containing laminin, whereas ductal and fibrotic compartments are increasingly exposed to collagen- and fibronectin-associated cues [4,16,17]. A reductionist hydrogel system therefore offers a useful way to control substrate mechanics independently from adhesive ligand composition. PAA hydrogels are particularly suited to this purpose because their elastic modulus can be tuned over a broad range by altering acrylamide and bis-acrylamide content while the surface is subsequently functionalized with defined adhesive ligands [18]. Previous work has also established robust workflows for mechanical characterization of compliant hydrogels by nanoindentation [19].

The protocol presented here establishes three PAA stiffness conditions spanning a soft, medium, and fibrotic-like range and combines these with fibronectin-derived arginine-glycine-aspartate (RGD) peptide or a mixed RGD/laminin-derived isoleucine-lysine-valine-alanine-valine (IKVAV) ligand condition. Murine pancreatic acinar 266-6 cells are used as a reproducible screening model. Acinar identity is assessed by alpha-amylase 2 (AMY2), while YAP nuclear-to-cytoplasmic localization is quantified as a mechanotransduction readout. The method additionally incorporates a fluorescent RGD calibration step because identical soluble peptide inputs do not necessarily produce identical coupled ligand densities across PAA formulations. To demonstrate the flexibility of the same hydrogel chemistry, the platform is also implemented with gelatin as an alternative surface ligand at comparable soft and stiff mechanical endpoints, with the ductal-lineage marker sex-determining region Y-box transcription factor 9 (SOX9) and quantitative cell morphology used as complementary phenotypic readouts. These configurations collectively test whether matrix stiffness remains a dominant determinant of pancreatic acinar cell state across different ligand environments and readout strategies. The overall aim is therefore not only to separate stiffness from ECM composition, but also to provide an experimentally tractable platform in which the mechanical regime, surface ligand environment, and phenotypic readout can each be adapted to interrogate mechanosensitive features of ADM.

The overall experimental workflow, from tunable hydrogel fabrication and mechanical characterization through extracellular-matrix functionalization, pancreatic acinar cell culture, immunostaining, confocal imaging, and quantitative analysis, is summarized in **Figure 1**.

**Figure 1.**
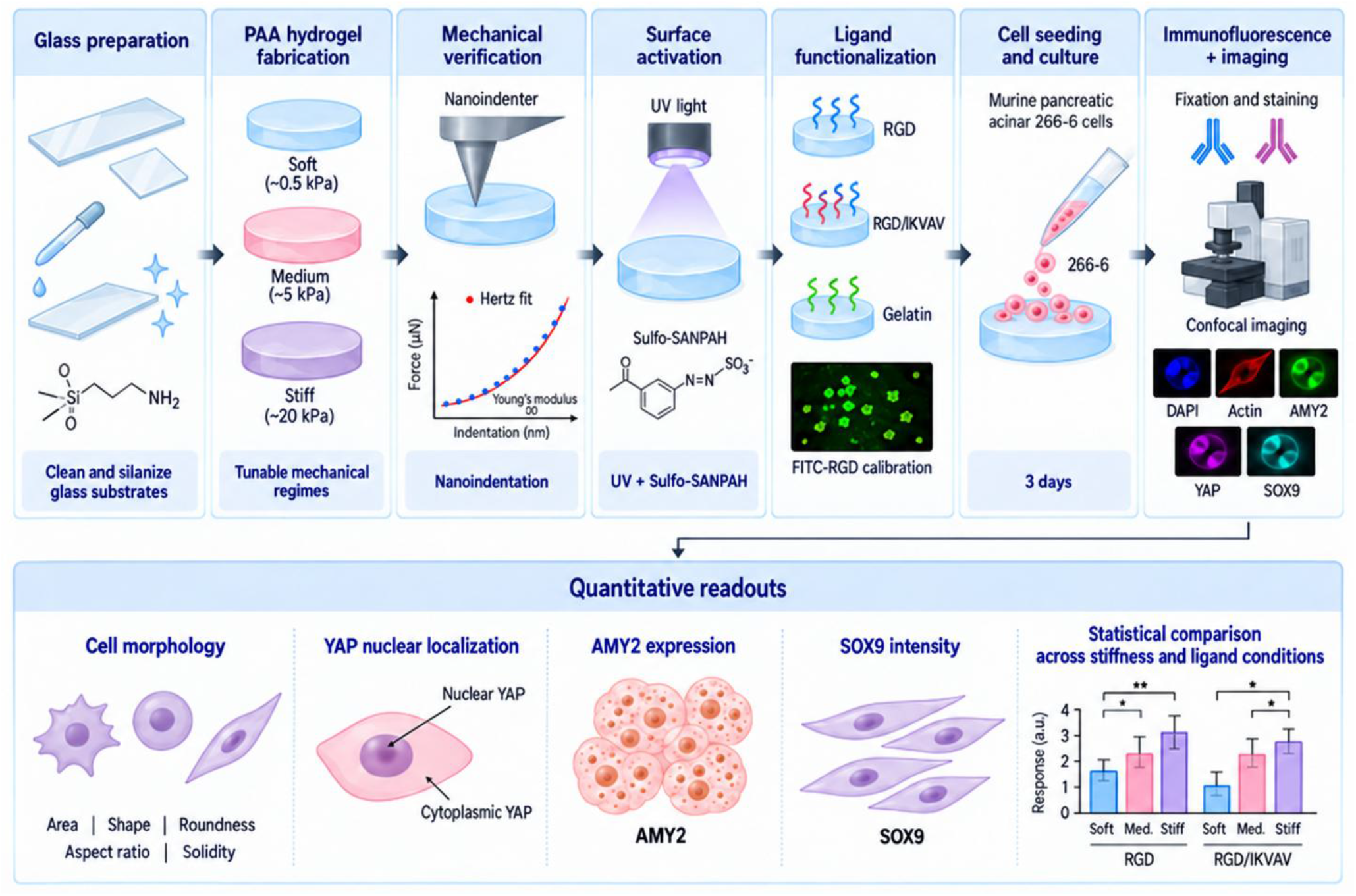
Experimental workflow for engineering tunable synthetic fibrotic niches to study acinar-to-ductal metaplasia in two-dimensional pancreatic culture systems. The workflow begins with preparation and silanization of glass substrates followed by fabrication of tunable PAA hydrogels spanning soft, medium, and stiff mechanical regimes. Hydrogel mechanical properties are verified by Chiaro nanoindentation, with force-indentation data fitted using the Hertz model to calculate Young’s modulus and confirm the intended stiffness range. The hydrogel surface is then activated using Sulfo-SANPAH under ultraviolet illumination and functionalized with extracellular-matrix ligands, including RGD, mixed RGD/IKVAV, or gelatin. Ligand coupling can be assessed using FITC-RGD as a calibration control. Murine pancreatic acinar 266-6 cells are seeded onto the functionalized hydrogel substrates and cultured for 3 days before fixation and immunofluorescence staining. ADM-associated and mechanotransductive responses are assessed by imaging AMY2, SOX9, YAP, filamentous actin, and nuclei by confocal microscopy. Quantitative image analysis is then used to assess cell morphology, YAP nuclear localization, AMY2 expression, and SOX9 intensity, followed by statistical comparison across hydrogel stiffness and ligand conditions. The schematic was generated using ChatGPT (OpenAI) based on an author-defined experimental workflow and was subsequently reviewed and verified by the authors for scientific accuracy.

## PROTOCOL

Ethics statement: This protocol uses an established murine pancreatic acinar cell line and does not involve live vertebrate animals, human participants, or primary human tissue. Refer to the Table of Materials for supplier, catalogue, equipment, and software details used throughout the protocol.

### 1. Prepare glass substrates

1.1 Use 12 mm glass coverslips and glass microscope slides for hydrogel fabrication. Sonicate both in ultrapure water for 30 min, transfer them to ethanol, and sonicate for a further 30 min.
1.2 Rinse the cleaned glass with ultrapure water and dry it under a stream of nitrogen. Keep the cleaned slides and coverslips free of dust and handle them with clean forceps.
1.3 CAUTION: Perform silanization procedures in a chemical fume hood and follow the relevant safety data sheets and institutional chemical-safety procedures. Passivate each glass slide by immersing it in a hydrophobic silanizing reagent for less than 5 s, allow it to air-dry, and then sonicate it in ethanol for 15 min.
1.4 Prepare the coverslip silanization solution by combining 50 mL ethanol, 2.5 mL ultrapure water, and 231 µL 3-(Trimethoxysilyl)propyl methacrylate. Immerse the coverslips for 120 min to create an acrylate-reactive surface that can covalently anchor the PAA gel [20].
1.5 Remove the coverslips from the silanization solution, dry them under nitrogen, and heat them at 120 °C for 60 min. Allow the coverslips to cool to room temperature before hydrogel fabrication.

### 2. Fabricate tunable polyacrylamide hydrogels

2.1 CAUTION: Unpolymerized acrylamide is hazardous. Prepare acrylamide-containing solutions with appropriate personal protective equipment, avoid skin contact, and dispose of unpolymerized waste according to institutional procedures.
2.2 Prepare the polyacrylamide (PAA) prepolymer solutions by combining the required volumes of acrylamide, N,N′-methylenebisacrylamide, and PBS to generate the desired soft, medium, and stiff hydrogel formulations. Adjust the relative acrylamide and bis-acrylamide content to achieve the intended mechanical regime. Refer to Table 1 for the specific component volumes used for each 1 mL hydrogel formulation [18].
2.3 Degas each acrylamide/bis-acrylamide/PBS mixture in a desiccator for 20 min to reduce dissolved oxygen, which inhibits free-radical polymerization. Immediately before casting, add 1 µL tetramethylethylenediamine (TEMED) and 10 µL of 10% ammonium persulfate to each 1 mL formulation and mix gently without introducing bubbles.
2.4 Pipette 10 µL of activated prepolymer onto a passivated glass slide. Immediately lower an acrylsilanized coverslip onto the droplet with the treated surface facing the prepolymer, allowing the droplet to spread beneath the coverslip.
2.5 Allow the gel to polymerise undisturbed for approximately 30 min at room temperature, checked by monitoring solidification of a parallel gel mixture left in an Eppendorf tube. Once set, separate the coverslip carefully from the passivated slide so that the PAA gel remains covalently attached to the coverslip. NOTE: polymerisation time may vary slightly depending on room temperature; if coverslips are challenging to separate from glass slides, incrementally reduce polymerization time until separation can be achieved. Soaking polymerized gels in PBS for 30+ mins can help with detachment.
2.6 Place each gel-coated coverslip gel-side-up in a non-tissue-culture-treated 24-well plate containing PBS and allow the gel to swell overnight. PAUSE POINT: Swollen gels can be held in PBS before subsequent mechanical characterization and functionalization, provided sterility and hydration are maintained.

**Table 1.** Polyacrylamide master-mix formulations used to generate soft, medium, and stiff hydrogels. Volumes of 40% acrylamide, 2% N,N’-methylenebisacrylamide, PBS, TEMED, and 10% ammonium persulfate are provided for a final volume of 1 mL per formulation.

| Sample | 40%<br>acrylamide<br>(μL) | 2%<br>N,N'-methylenebisacrylamide<br>(μL) | PBS<br>(μL) | TEMED<br>(μL) | 10%<br>ammonium<br>persulfate<br>(μL) |
| --- | --- | --- | --- | --- | --- |
| Soft (0.5<br>kPa) | 100 | 15 | 874 | 1 | 10 |
| Medium<br>(4 kPa) | 100 | 75 | 814 | 1 | 10 |
| Stiff (18<br>kPa) | 250 | 50 | 689 | 1 | 10 |

### 3. Verify hydrogel mechanics by nanoindentation

3.1 Equilibrate the hydrogel samples in PBS before testing and mount the gel-coated coverslip securely on the nanoindentation stage. Calibrate the probe according to the instrument workflow before contacting the sample surface [19].
3.2 Perform nanoindentation using the Chiaro nanoindenter fitted with probe P250745M. Use a probe spring constant of 0.45 N/m and a spherical tip radius of 28 µm. Set the loading rate to 5 µm/s and the maximum indentation depth to approximately 5 µm.
3.3 Acquire 25 indentation measurements at spatially separated positions on each hydrogel and collect force–distance curves from three technical hydrogel replicates per stiffness condition, yielding a maximum of 75 curves per condition. Avoid the gel edge and visibly damaged, irregular, or dehydrated regions.
3.4 Analyze the force–distance curves using DataViewer v3.4.7. Identify the contact point and convert the force–distance data to force–indentation curves. Fit the post-contact indentation region, extending from the identified contact point to the maximum indentation depth of approximately 5 µm, using a Hertz contact mechanics model to determine Young’s modulus. Assume a Poisson’s ratio of 0.5 for all hydrogel samples and apply the same fitting assumptions and acquisition settings across all stiffness conditions.
3.5 For quality control, retain only curves with a clearly defined baseline and contact point and with several micrometres of measurable indentation into the hydrogel. Exclude curves with an unsuitable baseline, poorly defined contact point, or insufficient measurable indentation from subsequent analysis.
3.6 Confirm that the measured Young’s moduli fall within the intended stiffness ranges before proceeding with cell experiments. In the representative dataset, mean Young’s moduli were 0.50 ± 0.21 kPa, 4.48 ± 0.15 kPa, and 18.29 ± 0.88 kPa for the soft, medium, and stiff hydrogel conditions, respectively.

### 4. Functionalize hydrogels with ECM-derived ligands

4.1 Prepare a 0.2 mg/mL solution of the photoactivatable crosslinker sulfosuccinimidyl 6-(4′-azido-2′-nitrophenylamino)hexanoate (Sulfo-SANPAH) in ultrapure water immediately before use. Protect the solution from light until exposure.
4.2 CAUTION: Use appropriate shielding and eye/skin protection for ultraviolet illumination. Cover each hydrated PAA surface with the crosslinker solution and expose it to 324 nm ultraviolet light for 10 min using the ultraviolet illumination system.
4.3 Remove the crosslinker solution and wash the activated gels three times with 50 mM 4-(2-hydroxyethyl)-1-piperazineethanesulfonic acid (HEPES) buffer, adjusted to pH 8.5, for 5 min per wash. Ensure that the gels remain fully submerged in HEPES buffer throughout each wash to remove unreacted material.
4.4 Prepare ligand solutions for the relevant experimental configuration. For the defined peptide series, prepare RGD-only and mixed RGD/IKVAV conditions, using 80% RGD and 20% IKVAV by molar amount for the mixed condition. To demonstrate adaptation to a protein ligand, prepare 0.1% (w/v) bovine-skin gelatin; the representative gelatin-functionalized configuration used the soft and stiff mechanical endpoints.
4.5 Add the appropriate RGD, RGD/IKVAV, or gelatin ligand solution to each activated hydrogel and incubate at 37 °C overnight to couple ligand amines to the activated surface. PAUSE POINT: Complete the overnight coupling before cell seeding on the following day.
4.6 For the defined peptide series, use a nominal total peptide concentration of 500 µM for the attachment-optimized condition. Because coupling efficiency differs across PAA formulations, determine and apply a stiffness-specific correction if equal effective ligand density is required; the representative dataset used a relative correction factor of 1.63:1.00:0.77 for soft:medium:stiff gels.
4.7 For all functionalized hydrogels, remove excess non-coupled peptide/protein after the overnight incubation by washing three times with PBS for 5 min per wash, using 500 µL per well in a 24-well plate.

### 5. Calibrate peptide coupling and optimize cell attachment

5.1 To quantify stiffness-dependent peptide coupling, prepare RGD solutions containing 1% FITC-labelled RGD and 99% unlabeled RGD. Include a gel without FITC-RGD as a negative fluorescence control.
5.2 Image three spatially separated regions from each of three technical hydrogel replicates per stiffness condition by confocal fluorescence microscopy. Use identical acquisition settings within each calibration experiment and avoid saturation.
5.3 Subtract the negative-control background and quantify mean fluorescence intensity in ImageJ. Calculate a stiffness-specific concentration correction relative to the medium condition and repeat the fluorescence measurement after correction to verify that the systematic stiffness-dependent gradient has been reduced.
5.4 For attachment optimization, functionalize gels with RGD at 50, 250, or 500 µM and seed cells as described in section 6. After 1 day, fix and stain the cultures with phalloidin and 4′,6-diamidino-2-phenylindole (DAPI), then assess cell number, clustering, and spread morphology.
5.5 Select the concentration that provides robust attachment while preserving resolvable individual cells and small clusters. In the representative dataset, 500 µM RGD produced the best overall attachment and spreading across the three stiffnesses.

### 6. Expand and seed pancreatic acinar cells

6.1 Thaw cryopreserved murine pancreatic acinar 266-6 cells and expand them in 75 cm^2^ (T75) tissue-culture flasks. The representative experiments used cells cryopreserved at passage 2 and expanded to passage 3, which was maintained for the study.
6.2 Culture the cells in high glucose Dulbecco’s Modified Eagle’s Medium (DMEM) supplemented with 10% fetal bovine serum, 1% antibiotic-antimycotic solution, and 1% non-essential amino-acid supplement.
6.3 Allow the cells to expand for approximately 48 h before seeding. Maintain the same passage range and culture history across stiffness and ligand conditions to reduce cell-state variability.
6.4 Detach the cells using 1× trypsin-EDTA dissociation reagent and incubate for 5 min at 37 °C in the tissue-culture incubator. Collect the detached cells, centrifuge at 300 × g, remove the dissociation reagent, and resuspend the cell pellet in complete culture medium to terminate dissociation. Count the cells using the haemocytometer and seed 50,000 cells onto each functionalized hydrogel in a final volume of 500 µL per well. Before cell seeding, maintain the hydrogels in PBS and expose them to the tissue-culture hood ultraviolet lamp for 30 min for sterilization.
6.5 For the defined peptide dataset, prepare the full matrix of nine conditions: three stiffnesses (soft, medium, stiff) crossed with RGD and RGD/IKVAV surfaces. Use three technical hydrogel replicates per condition for YAP staining and a separate three technical hydrogel replicates per condition for AMY2 staining. To demonstrate ligand and readout flexibility within the same platform, prepare soft and stiff gelatin-functionalized hydrogels using three technical hydrogel replicates per condition for SOX9 and morphology analysis.
6.6 Culture the cells on the functionalized hydrogels for 3 days at 37 °C in a humidified incubator with 5% CO_2_. Maintain a final culture-medium volume of 500 µL per well and replace the culture medium at 3-day intervals.

### 7. Fix and immunostain for YAP, AMY2, or SOX9

7.1 At day 3, aspirate the culture medium and fix the cells with 4% formaldehyde for 15 min. Wash the samples three times with PBS for 5 min per wash.
7.2 Permeabilize the cells with 0.1% Triton X-100 for 5 min at room temperature and wash three times with PBS for 5 min per wash. Block nonspecific binding with 1% bovine serum albumin (BSA) prepared in PBS for 30 min.
7.3 For YAP or AMY2 staining, dilute the corresponding primary antibody to 1:450 in blocking solution. For SOX9 staining on gelatin-functionalized hydrogels, dilute the SOX9 primary antibody to 1:250 in blocking solution. Incubate primary antibodies overnight at 4 °C and use separate hydrogel samples for each marker workflow.
7.4 The following day, wash the samples three times with 0.05% Tween-20 in PBS for 5 min per wash, using 500 µL per wash, before secondary labeling.
7.5 For YAP and AMY2 staining, incubate the samples for 1 h with a Cy3-conjugated anti-rabbit secondary antibody diluted 1:400. For SOX9 staining, use a Cy3-conjugated anti-mouse secondary antibody at 1:500. In both workflows, co-incubate with phalloidin diluted 1:600 in blocking solution. Protect the samples from light during and after secondary-antibody incubation.
7.6 Wash the samples three times with 0.05% Tween-20 in PBS for 5 min per wash, using 500 µL per wash, to remove unbound secondary antibody and phalloidin. Mount the coverslips on glass slides using a DAPI-containing mounting medium and allow the mounted samples to cure for at least 1 h before imaging.

### 8. Acquire confocal images

8.1 Acquire confocal z-stacks using an LSM 980 confocal microscope equipped with a Plan-Apochromat 20×/0.8 M27 air objective. Acquire DAPI, phalloidin/AF488, and Cy3-labelled marker channels using 405, 488, and 561 nm excitation laser lines, respectively. Use separate acquisition tracks for each fluorescence channel to minimize spectral overlap.
8.2 Acquire images at 512 × 512 pixels with an XY pixel size of 0.637 × 0.637 µm, a z-step of 0.73 µm, and a scan zoom of 1.3×. Set the pinhole to approximately 1 Airy unit for each channel. Use detection windows of 410.01–535.86 nm for DAPI, 490.50–545.91 nm for AF488, and 530.05–757.11 nm for the Cy3/RhodB channel. Set detector gains to 500 V for DAPI and 700 V for both AF488 and Cy3/RhodB, with a digital gain of 1 and digital offset of 0 for all channels. Acquire images at 16-bit depth using bidirectional scanning and mean line averaging of 1×.
8.3 Record sufficient optical sections to encompass the full cell height so that subsequent maximum-intensity projection and region masking represent the entire cell. A representative z-stack comprised 30 optical sections.
8.4 Acquire at least three spatially separated fields of view per hydrogel from three technical hydrogel replicates for each experimental condition. Use a consistent sampling strategy across all hydrogel conditions and avoid visibly damaged, dehydrated, or otherwise unrepresentative regions.
8.5 Use identical acquisition settings for all conditions compared within an experiment and ensure that no fluorescence channel is saturated. NOTE: Imaging in the representative study was not performed blinded to experimental condition; the operator was aware of sample identity during acquisition.

### 9. Quantify YAP localization, AMY2 and SOX9 expression, and cell morphology in ImageJ

9.1 Open each confocal z-stack in ImageJ v1.54p and generate a maximum-intensity projection. Keep the DAPI, phalloidin, and marker channels registered and use the same preprocessing steps for all images.
9.2 For nuclear regions, threshold the DAPI channel and convert it to a binary mask. Invert the mask if required by the analysis workflow and divide pixel values by 255 to create a normalized binary mask containing values of 0 and 1.
9.3 Multiply the normalized nuclear mask by the YAP channel using Image Calculator to isolate nuclear YAP signal. Measure the nuclear area and raw integrated density (RawIntDen), then calculate mean nuclear YAP intensity as RawIntDen/area.
9.4 Threshold the phalloidin channel to create a whole-cell mask and repeat the masking procedure to isolate whole-cell YAP signal. Calculate cytoplasmic area and integrated intensity by subtracting the nuclear values from the corresponding whole-cell values.
9.5 Calculate mean cytoplasmic YAP intensity and then compute the nuclear-to-cytoplasmic ratio by dividing the mean nuclear intensity by the mean cytoplasmic intensity. Use this ratio as the quantitative measure of YAP nuclear localization.
9.6 For AMY2, use the phalloidin-derived whole-cell mask to isolate the AMY2 channel and calculate mean AMY2 intensity as RawIntDen/area. Normalize each individual AMY2 value to the mean of the soft RGD reference condition so that the reference mean equals 1.
9.7 Manually threshold the DAPI and phalloidin channels for each image so that distinct cellular staining is clearly separated from background signal while accounting for image-to-image variation. Exclude cells that are not fully contained within the field of view and exclude fluorescence signal that does not spatially correspond with the expected nuclear or cytoskeletal compartments defined by DAPI and phalloidin, respectively. Generate ROIs on a per-cell basis so that each cell is quantified separately and can be tracked consistently throughout the analysis workflow.
9.8 For SOX9 quantification, manually threshold the phalloidin channel to delineate the complete cellular area and convert the thresholded image to an 8-bit binary mask. Divide the binary image by 255 using Process > Math > Divide to generate a mask in which background pixels have a value of 0 and cell-associated pixels have a value of 1. Using Process > Image Calculator, multiply the resulting 0/1 cell mask by the corresponding SOX9 fluorescence image to retain SOX9 signal only within the phalloidin-defined cellular area. Generate per-cell ROIs and normalize the measured SOX9 fluorescence signal to the phalloidin-defined cell area for each analyzed cell.
9.9 For morphology analysis, quantify roundness, aspect ratio, and geometric regularity/solidity from the phalloidin-defined cell boundary. Calculate geometric regularity/solidity as the ratio of segmented cell area to convex-hull area (Area/Convex Area). Values approaching 1 indicate a more convex and geometrically regular cell outline, whereas lower values indicate a more irregular or indented cell shape.

### 10. Statistical analysis and quality control

10.1 Perform all statistical analyses using Prism v10.5.0. Report the number of technical hydrogel replicates (N) separately from pooled observations, such as analyzed cells, fields of view, or indentation curves (n). Pool measurements obtained from technical hydrogel replicates within each experimental condition for statistical analysis, such that n represents the total number of pooled observations and N represents the number of technical hydrogel replicates.
10.2 Apply the relevant experimental quality-control criteria before statistical analysis and exclude measurements that do not meet the predefined acceptance criteria. Identify potential statistical outliers using the robust regression and outlier removal (ROUT) method with Q = 1%.
10.3 Evaluate the statistical assumptions for each dataset before selecting the appropriate test. Use descriptive statistics to assess the variability between groups. Assume equal standard deviations only when the standard deviations are comparable; if the largest group standard deviation is more than twofold greater than the smallest, do not assume equal standard deviations. Assess data distribution using the D’Agostino–Pearson, Anderson–Darling, Shapiro–Wilk, and Kolmogorov–Smirnov normality tests.
10.4 Select the appropriate parametric or nonparametric statistical analysis for each dataset on the basis of the normality and variance assessments described above. Report the specific statistical test, sample size, exact *P* value where practical, and the measure of central tendency and error representation in the corresponding figure or table legend.

## REPRESENTATIVE RESULTS

The hydrogel formulations produced three mechanically distinct PAA substrates spanning the intended physiological-to-fibrotic range. Nanoindentation yielded mean Young’s moduli of 0.50 ± 0.21 kPa for the soft condition, 4.48 ± 0.15 kPa for the medium condition, and 18.29 ± 0.88 kPa for the stiff condition. All pairwise comparisons were significant (P < 0.0001), demonstrating that the fabrication workflow can reproducibly separate the three mechanical environments (**Figure 2**).

**Figure 2.**
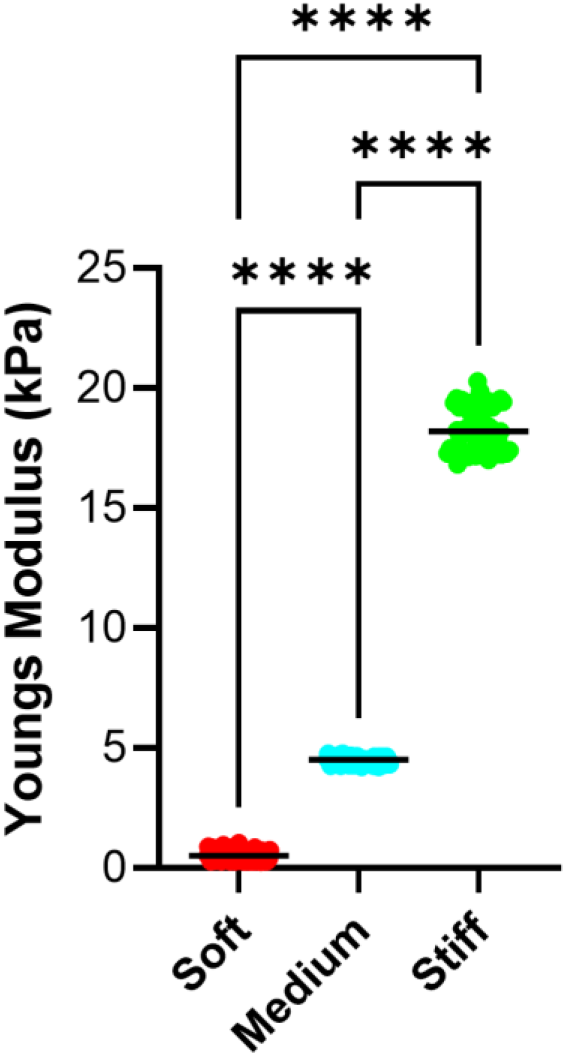
Mechanical characterization of the three PAA hydrogel formulations by nanoindentation. Mean Young’s modulus was 0.50 ± 0.21 kPa for soft gels, 4.48 ± 0.15 kPa for medium gels, and 18.29 ± 0.88 kPa for stiff gels. Individual plotted observations represent accepted force–distance curves pooled across N = 3 technical hydrogel replicates, with n = 73–75 curves retained per condition after quality-control exclusion. Data are presented as mean ± SD. All pairwise comparisons were significant using a Kruskal–Wallis analysis with Dunn’s multiple comparisons test (****P ≤ 0.0001).

An important quality-control observation was that peptide coupling varied with substrate stiffness when the same soluble RGD input concentration was used. Before correction, relative FITC-RGD intensity increased progressively from approximately 1.0 on soft gels to 2.1 on medium gels and 2.6 on stiff gels, corresponding to roughly a twofold increase from soft to medium and a 2.6-fold increase from soft to stiff. A stiffness-specific peptide-input correction factor of 1.63:1.00:0.77 for soft:medium:stiff gels was therefore applied. Following correction, the original monotonic stiffness-dependent increase was disrupted: relative FITC-RGD intensity was approximately 1.0, 1.3, and 0.7 for soft, medium, and stiff gels, respectively. Thus, the corrected conditions showed substantially reduced stiffness-dependent variation, although the medium condition remained significantly higher than the stiff condition (**Figure 3**).

**Figure 3.**
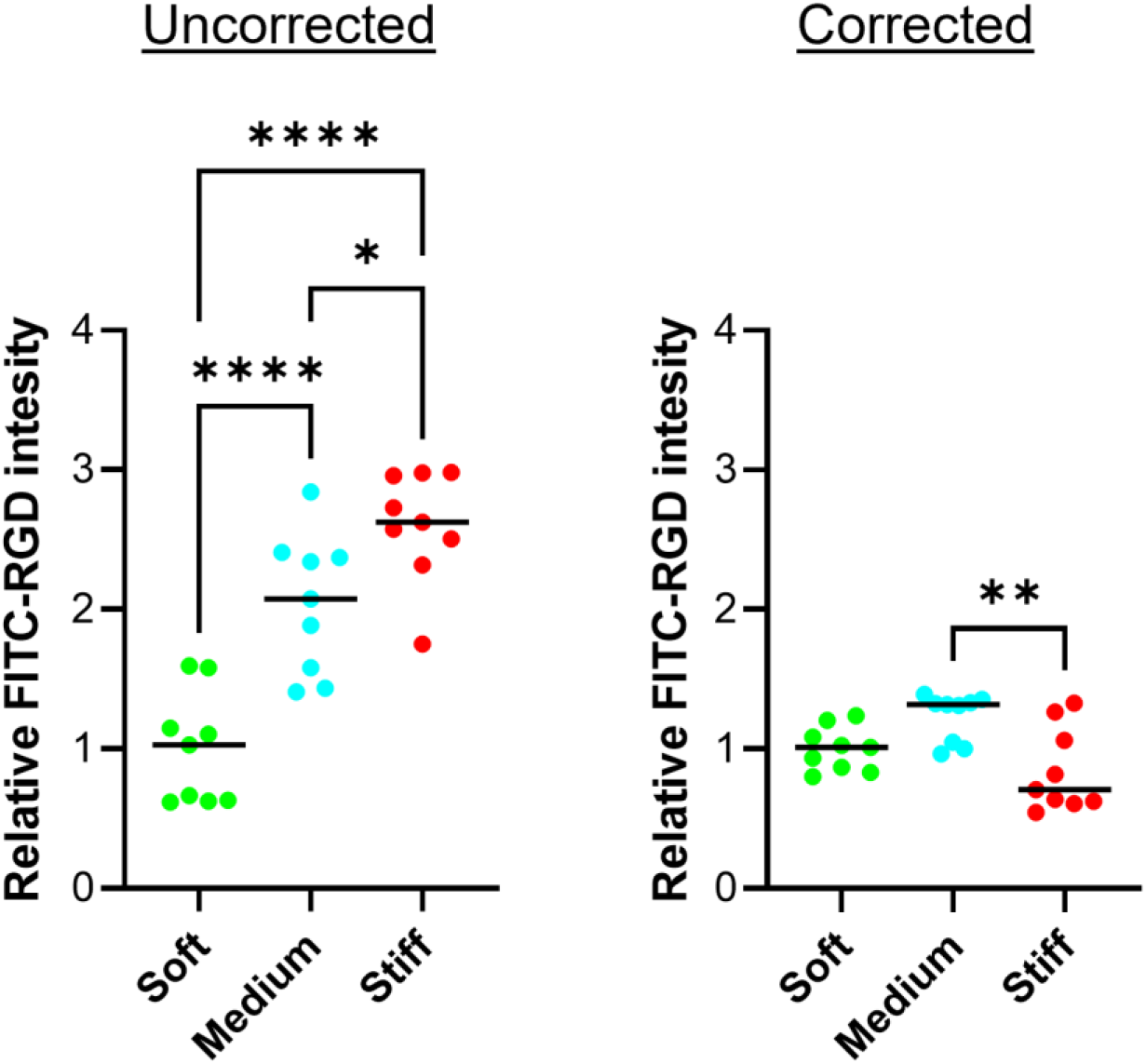
Quantification of FITC-RGD fluorescence before and after peptide-input correction. Before correction, fluorescence increased significantly with stiffness. After correction, the monotonic stiffness-dependent increase was disrupted, although a residual difference remained between conditions. Individual points represent imaged regions pooled across N = 3 technical hydrogel replicates per stiffness condition, with n = 9 imaged regions per condition. Data are presented as mean ± SD. Statistical significance was assessed using ordinary one-way ANOVA with Tukey’s multiple comparisons test; *P ≤ 0.05, **P < 0.01, ****P ≤ 0.0001 for the indicated comparisons.

Cell attachment and clustering after 1 day were influenced by both RGD concentration and substrate stiffness (**Table 2**). On soft hydrogels, 50 µM RGD produced substantial cell clustering, with an average of 12.2 cells per object and only 20% of objects representing single cells. Increasing the RGD concentration to 250 and 500 µM reduced the average number of cells per object to 3.9 and 2.4, respectively, while increasing the proportion of single cells to 50% and 60%. A similar trend was observed on medium-stiffness hydrogels, where the proportion of single cells increased from 33.3% at 50 µM RGD to 69.2% at 500 µM, with the average number of cells per object decreasing to 1.5 at the highest concentration. In contrast, cells on stiff hydrogels were predominantly present as isolated cells across all RGD concentrations tested, with an average of one cell per object and 100% single-cell objects. Mean object area generally decreased with reduced clustering on the soft substrates, whereas values were more comparable across RGD concentrations on medium and stiff substrates.

**Table 2.**
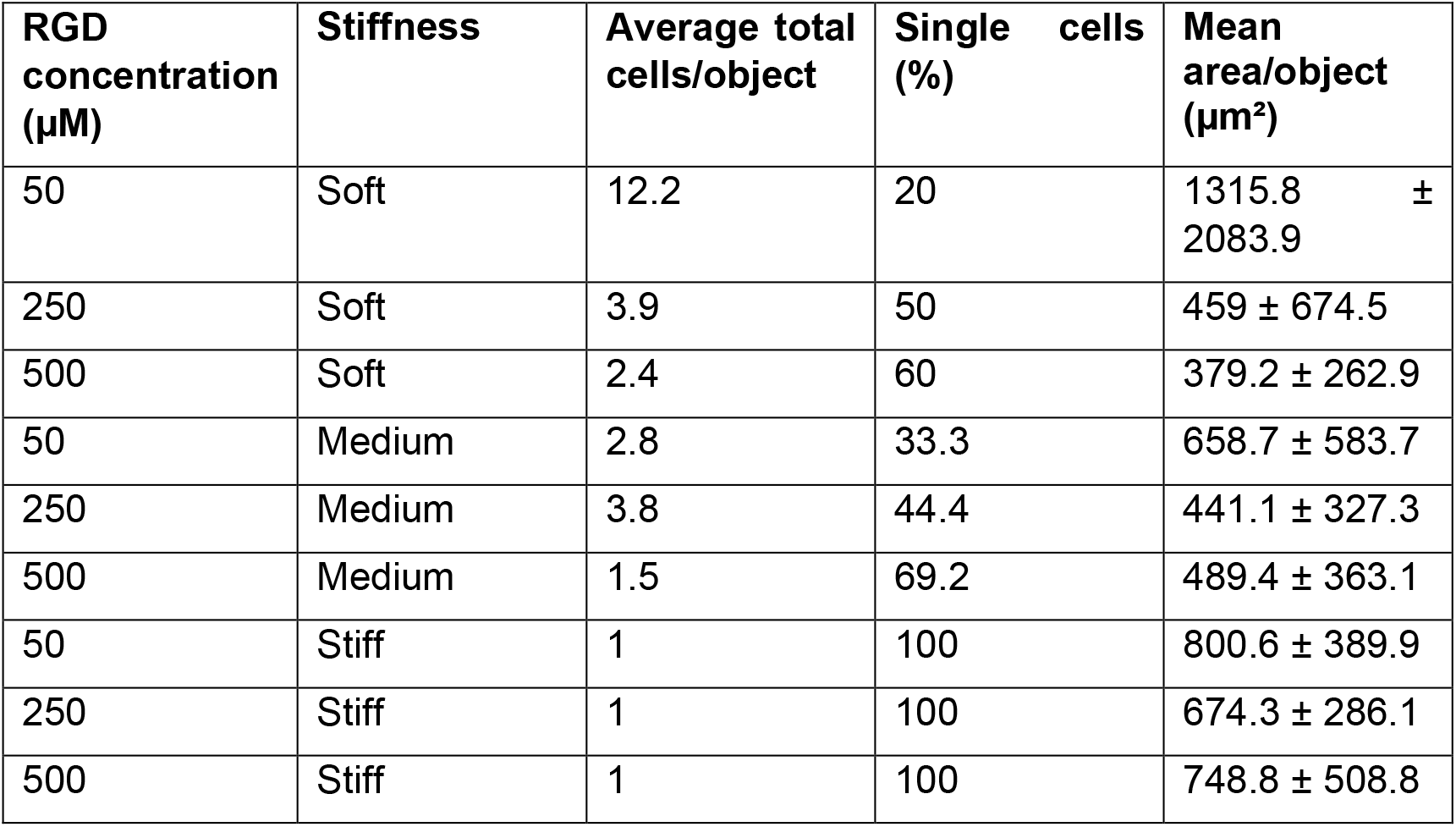
Cell attachment and clustering characteristics after 1 day across 50, 250, and 500 µM RGD and the three hydrogel stiffnesses. The table reports the average total cells per segmented object, percentage of single-cell objects, and mean object area for each condition. Where applicable, data are presented as mean ± SD; n = 6–14 segmented objects per condition.

Within the context of the phenotypic data obtained later in the study, this pattern is consistent with substrate mechanics influencing the balance between compact, acinar-associated organization and a more spread, ductal-associated phenotype. The greater clustering observed on soft substrates, particularly at lower RGD concentrations, is compatible with retention of a more compact acinar-like organization, whereas the predominance of individually resolved cells on stiff substrates is consistent with the increased spreading and elongation associated with the ductal-like phenotype observed at later time points. The progressive reduction in clustering with increasing RGD concentration on soft and medium gels also indicates that adhesive ligand availability can modulate cell organization within a given mechanical environment. However, these day-1 attachment measurements should be interpreted as an early morphological response rather than a direct measure of ADM. Together, the results support selection of 500 µM RGD for subsequent analyses because it provided robust attachment and individually resolvable cells while preserving the ability to distinguish stiffness-dependent phenotypic responses. At 500 µM RGD, single-cell morphology also varied with substrate stiffness: cell area and aspect ratio increased and circularity decreased between soft and stiff gels, while solidity followed a distinct stiffness-dependent pattern (**Figure 4**).

**Figure 4.**
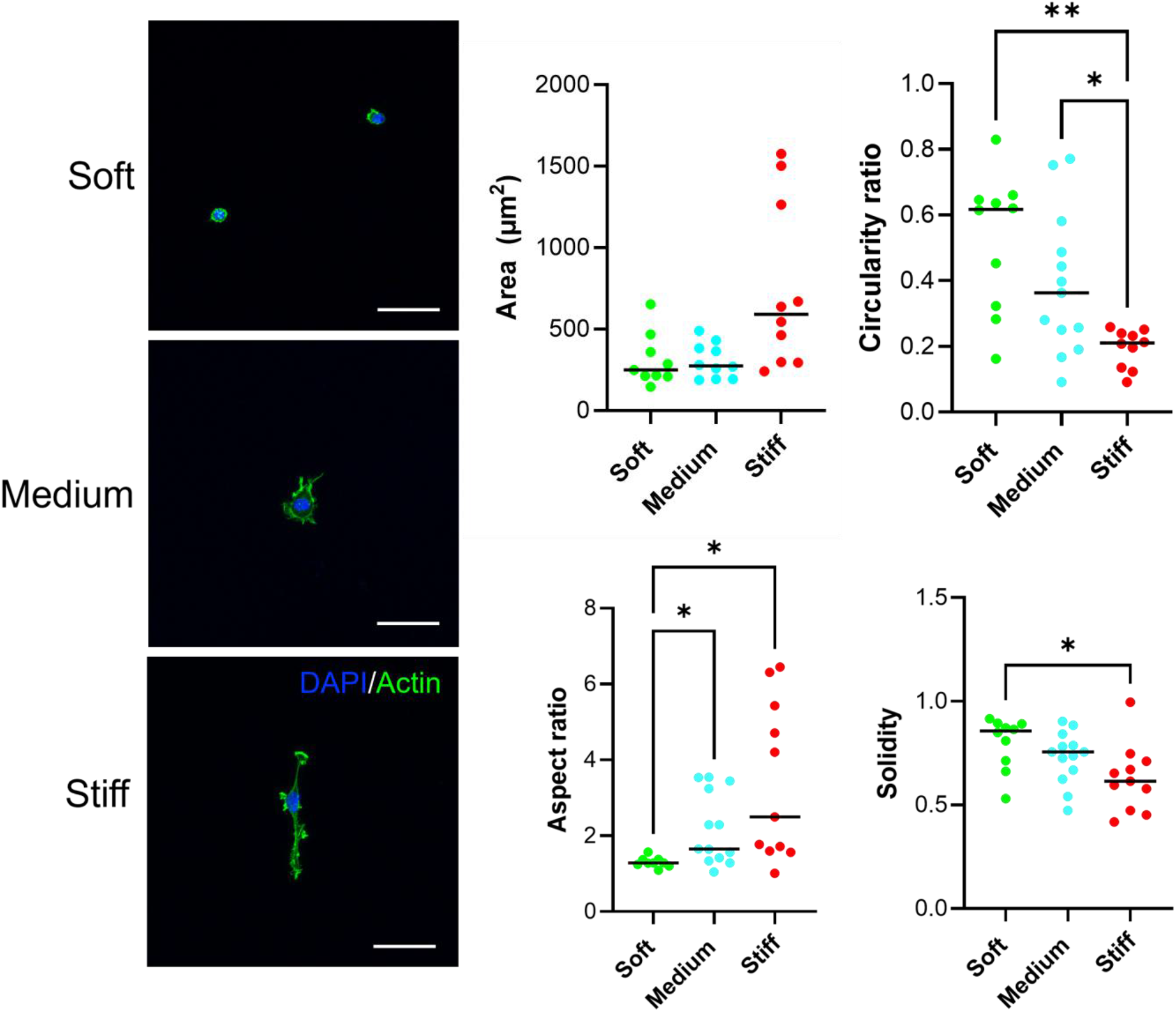
Single-cell morphology at 500 µM RGD as a function of substrate stiffness. Cell area, circularity, aspect ratio, and solidity were quantified from phalloidin-defined cell boundaries. Area and aspect ratio increased and circularity decreased between soft and stiff gels, while solidity followed a distinct stiffness-dependent pattern. Data are presented as mean ± SD; n = 10–13 analyzed cells per group. Statistical significance was assessed using ordinary one-way ANOVA or Brown–Forsythe and Welch ANOVA with Tukey’s and Dunnett’s T3 multiple comparisons tests; *P ≤ 0.05, **P ≤ 0.01. Scale bar = 30 µm.

AMY2 expression decreased as substrate stiffness increased. Within the RGD condition, normalized AMY2 fell from 1.00 on soft gels to 0.57 on stiff gels. The corresponding decrease in the RGD/IKVAV condition, from 0.85 to 0.66, did not remain significant. These AMY2 trends and representative immunofluorescence images are shown in **Figure 5**.

**Figure 5.**
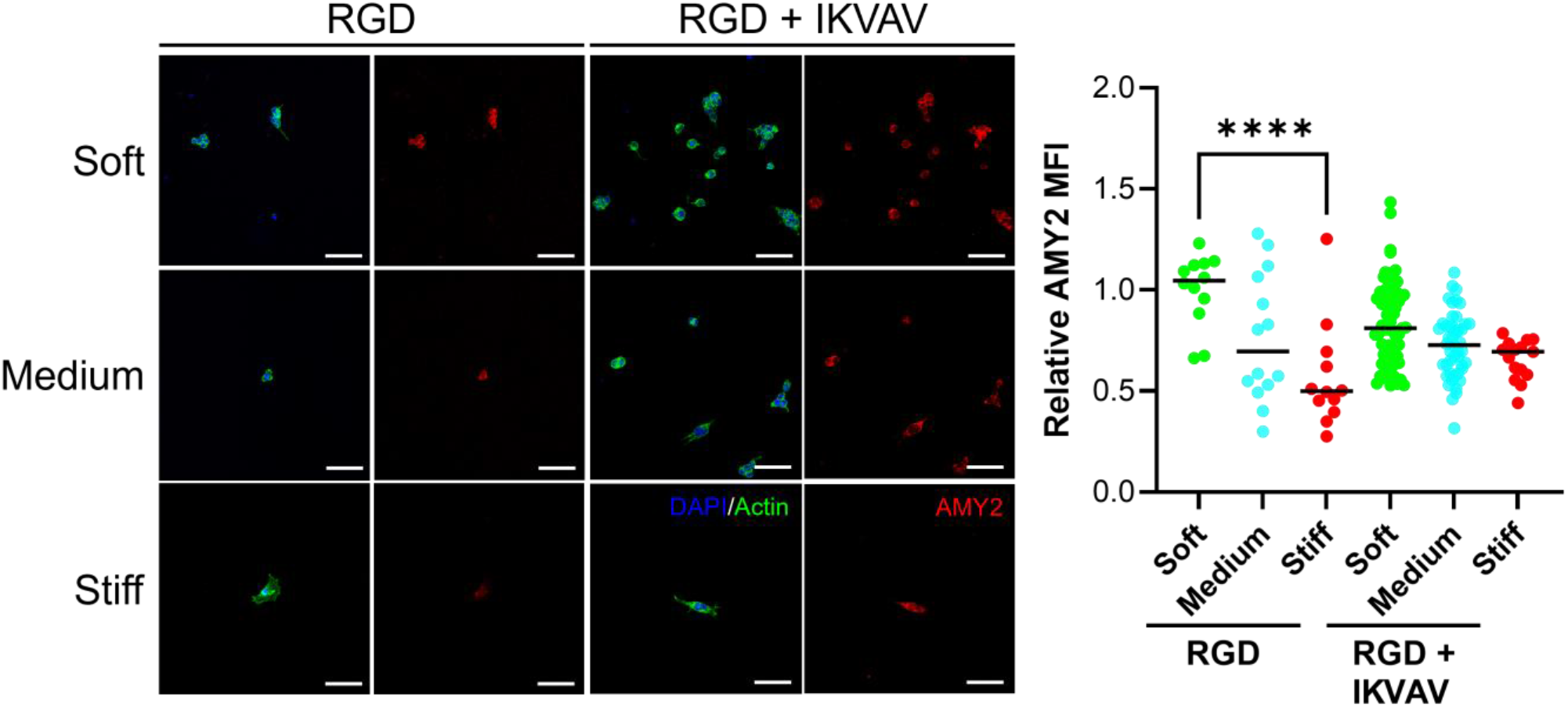
Relative AMY2 expression across stiffness and peptide conditions after 3 days. DAPI and phalloidin identify nuclei and cell morphology, respectively, and Cy3 identifies AMY2. Stiff RGD-only cells display a more spread phenotype and lower AMY2, whereas many cells on stiff RGD/IKVAV retain a more rounded phenotype. Whole-cell AMY2 intensity was normalized to the mean of the soft RGD condition. AMY2 decreased significantly from soft to stiff gels in the RGD condition, while the RGD/IKVAV soft-to-stiff comparison did not remain significant after correction for multiple comparisons. Cell-level measurements were pooled across N = 3 technical hydrogel replicates per condition; n = 10– 68 analyzed cells per group. Data are presented as mean ± SD. Statistical significance was assessed using a Kruskal–Wallis analysis with Dunn’s multiple comparisons test; ****P ≤ 0.0001 for the indicated comparisons. Scale bars = 30 µm.

YAP localization showed a complementary stiffness-dependent response. In RGD-functionalized gels, the nuclear-to-cytoplasmic (N:C) YAP ratio increased significantly from soft to medium and from soft to stiff. The mixed RGD/IKVAV condition showed the same pattern, with significant increases from soft to medium and soft to stiff. At any single stiffness, RGD and RGD/IKVAV did not differ significantly, supporting the conclusion that stiffness was the dominant determinant of YAP nuclear localization in the peptide-coupled conditions. Quantitative YAP localization data are shown in **Figure 6**.

**Figure 6.**
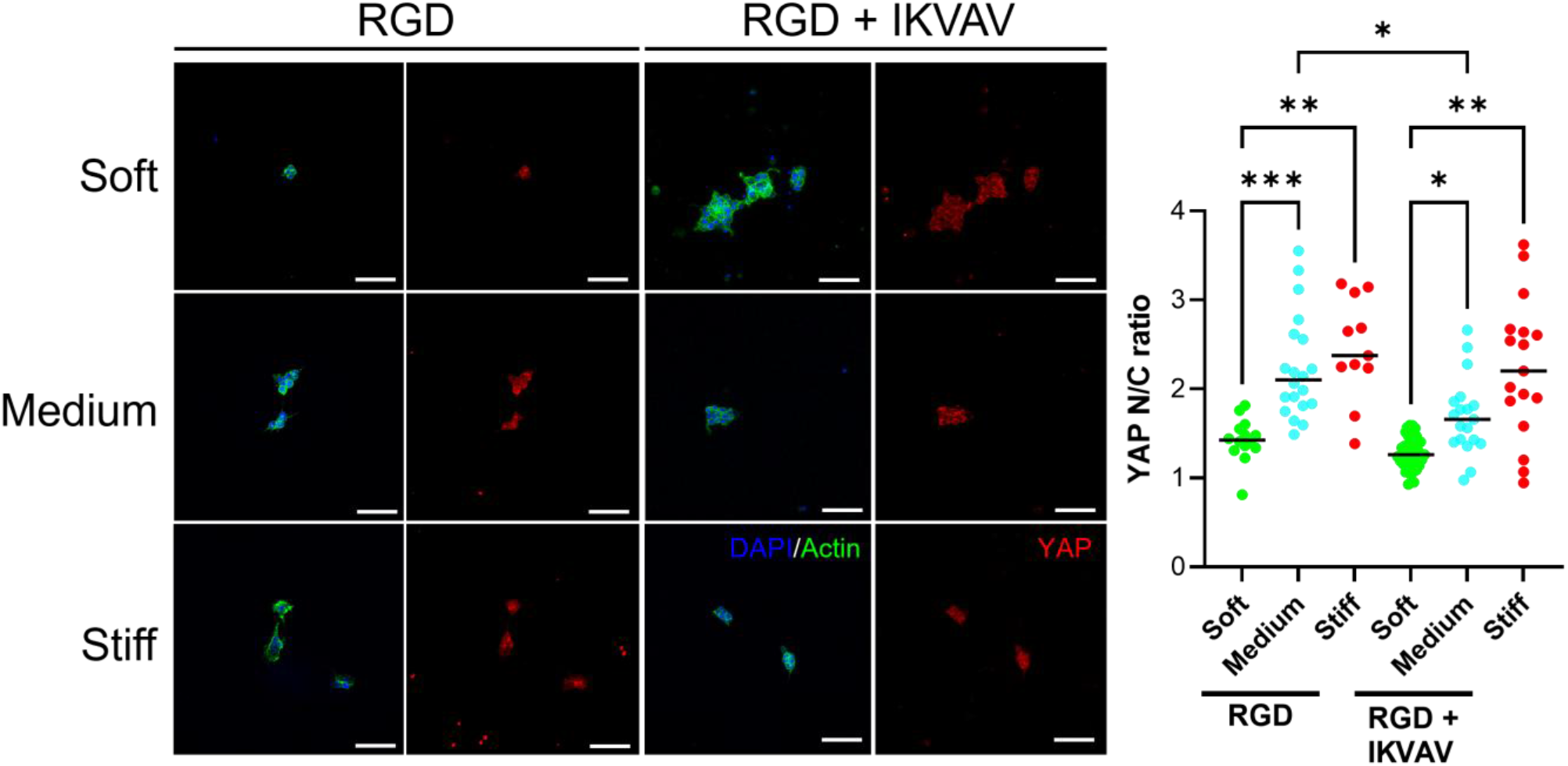
YAP nuclear localization and representative immunofluorescence across hydrogel stiffness and peptide conditions after 3 days. Representative YAP immunofluorescence images from RGD/IKVAV-functionalized soft, medium, and stiff gels, with a stiff RGD-only condition shown for comparison. DAPI and phalloidin identify nuclei and cell morphology, respectively, and Cy3 identifies YAP. YAP nuclear localization increases with substrate stiffness, while cells on stiff RGD/IKVAV gels retain a more acinar-like phenotype than cells on stiff RGD-only gels despite comparably high nuclear YAP. YAP localization was quantified as the nuclear-to-cytoplasmic fluorescence-intensity ratio. Significant increases from soft to medium and soft to stiff gels were detected in the RGD and RGD/IKVAV conditions. Cell-level measurements were pooled across N = 3 technical hydrogel replicates per condition; n = 11–38 analyzed cells per group. Data are presented as mean ± SD. Statistical significance was assessed using Brown–Forsythe and Welch ANOVA with Dunnett’s T3 multiple comparisons test; *P ≤ 0.05, **P ≤ 0.01, ***P ≤ 0.001 for the indicated comparisons. Scale bars = 30 µm.

Qualitative morphology provided additional information not captured by YAP or AMY2 intensity alone. Cells on soft gels retained compact acinar-like phenotype across peptide conditions. On stiff RGD-only gels, cells were more elongated and ductal-like and showed high nuclear YAP with reduced AMY2. On stiff RGD/IKVAV gels, however, many cells retained an acinar-like appearance despite similarly elevated nuclear YAP; the reported YAP N:C ratio in this condition was 2.23 ± 0.78. Thus, a successful experiment should reproduce a clear stiffness-dependent shift in YAP and AMY2 while also allowing the investigator to detect situations in which molecular and morphological readouts become dissociated. Representative images illustrating these morphology differences are included in **Figure 5 and 6**.

Extending the same PAA platform to gelatin as an alternative surface ligand and SOX9 as a complementary ductal-lineage readout further demonstrated the stiffness-dependent acinar-to-ductal phenotype. Murine pancreatic acinar 266-6 cells were cultured for 3 days on gelatin-presenting PAA hydrogels at soft and stiff endpoints. Compared with the soft condition, cells on the stiff substrate showed significantly reduced roundness, increased aspect ratio, and reduced geometric regularity/solidity, consistent with a shift from a compact acinar-like phenotype toward a more elongated phenotype associated with ductal marker expression. SOX9 fluorescence normalized to cell area was also markedly higher on the stiff substrate. All four soft-to-stiff comparisons were highly significant. Considered together with the AMY2 and YAP datasets, these results show a consistent stiffness-dependent cell-state shift across the adaptable configurations of the platform: softer matrices favor acinar-associated characteristics, whereas stiffer matrices favour an elongated phenotype associated with ductal marker expression. The use of gelatin alongside the defined RGD and RGD/IKVAV conditions further illustrates how ligand context and phenotypic readouts can be varied while maintaining matrix stiffness as the central mechanical input. The corresponding SOX9 imaging and quantification are shown in **Figure 7**.

**Figure 7.**
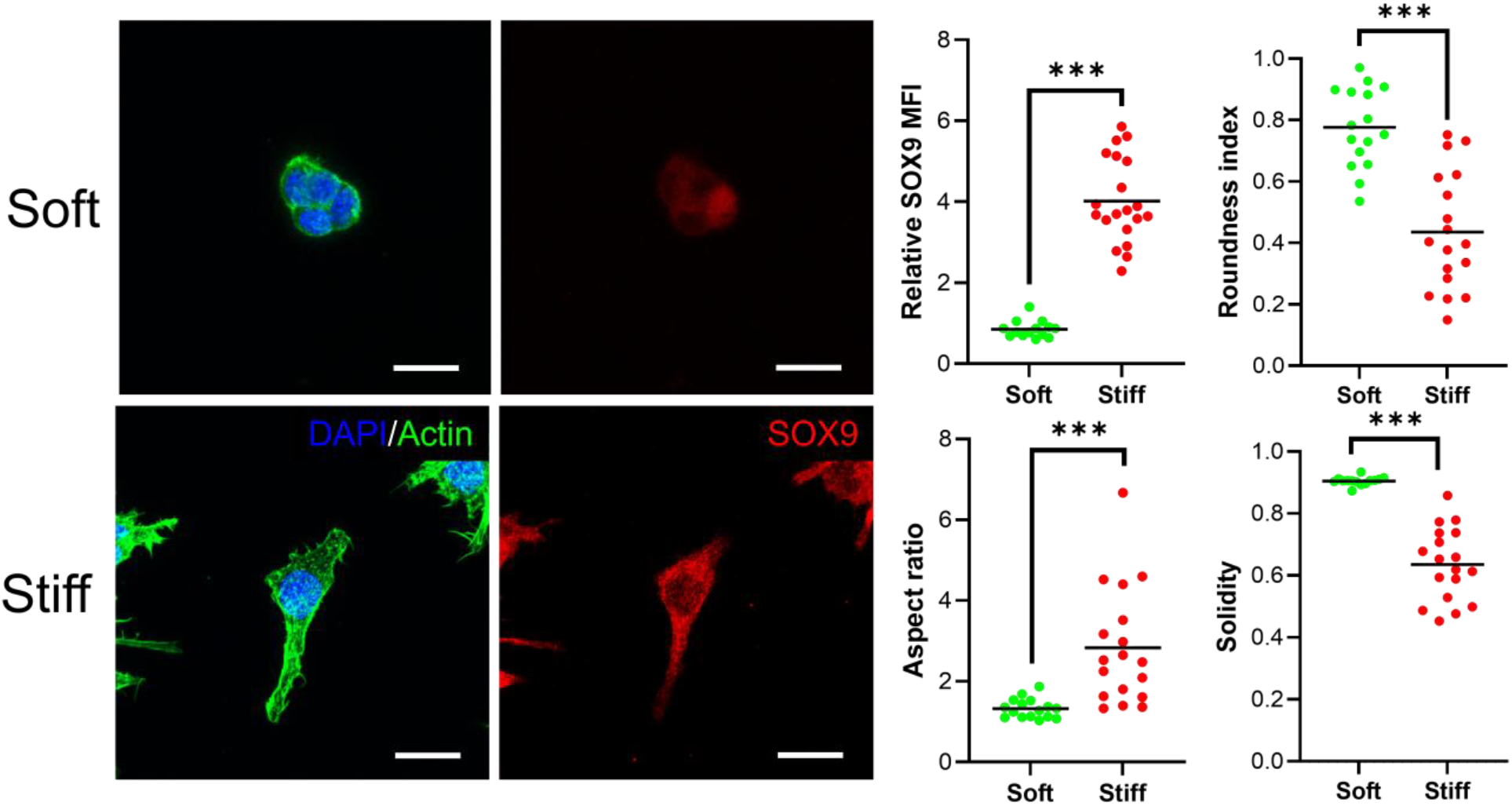
Substrate stiffness drives ductal-associated SOX9 expression in a gelatin-functionalized implementation of the PAA hydrogel platform. Murine pancreatic acinar 266-6 cells were cultured for 3 days on soft and stiff PAA hydrogels. Representative immunofluorescence images show nuclei (DAPI), actin (phalloidin), and SOX9; SOX9 fluorescence intensity normalized to cell area was markedly increased on stiff substrates. Cell-level measurements were pooled across N = 3 technical hydrogel replicates per condition; n = 16–18 analyzed cells per group. Data are presented as mean ± SD. The soft-versus-stiff comparison was significant by Welch’s t-test (***P ≤ 0.001). Scale bars = 10 µm.

Collectively, these findings demonstrate a stiffness-dependent progression from an acinar-associated phenotype on soft substrates toward an ADM-associated ductal phenotype on stiff substrates, with the medium mechanical condition representing a transitional cell state. Across the study, increasing stiffness was associated with progressive cell spreading and elongation, increased nuclear YAP localization, loss of AMY2 expression, and increased SOX9 expression in the soft-versus-stiff gelatin-functionalized configuration. Matrix stiffness was therefore the dominant determinant of cell phenotype across the ligand environments examined. Ligand composition nevertheless modulated how the mechanical response was expressed: cells on stiff RGD/IKVAV substrates retained a more acinar-like phenotype despite elevated nuclear YAP, indicating that laminin-derived IKVAV cues can partially buffer the phenotypic consequences of a high-stiffness environment without abolishing mechanotransductive signaling. The integrated relationship between matrix stiffness, ligand presentation, and ADM-associated phenotypic plasticity is summarized in **Figure 8**.

**Figure 8.**
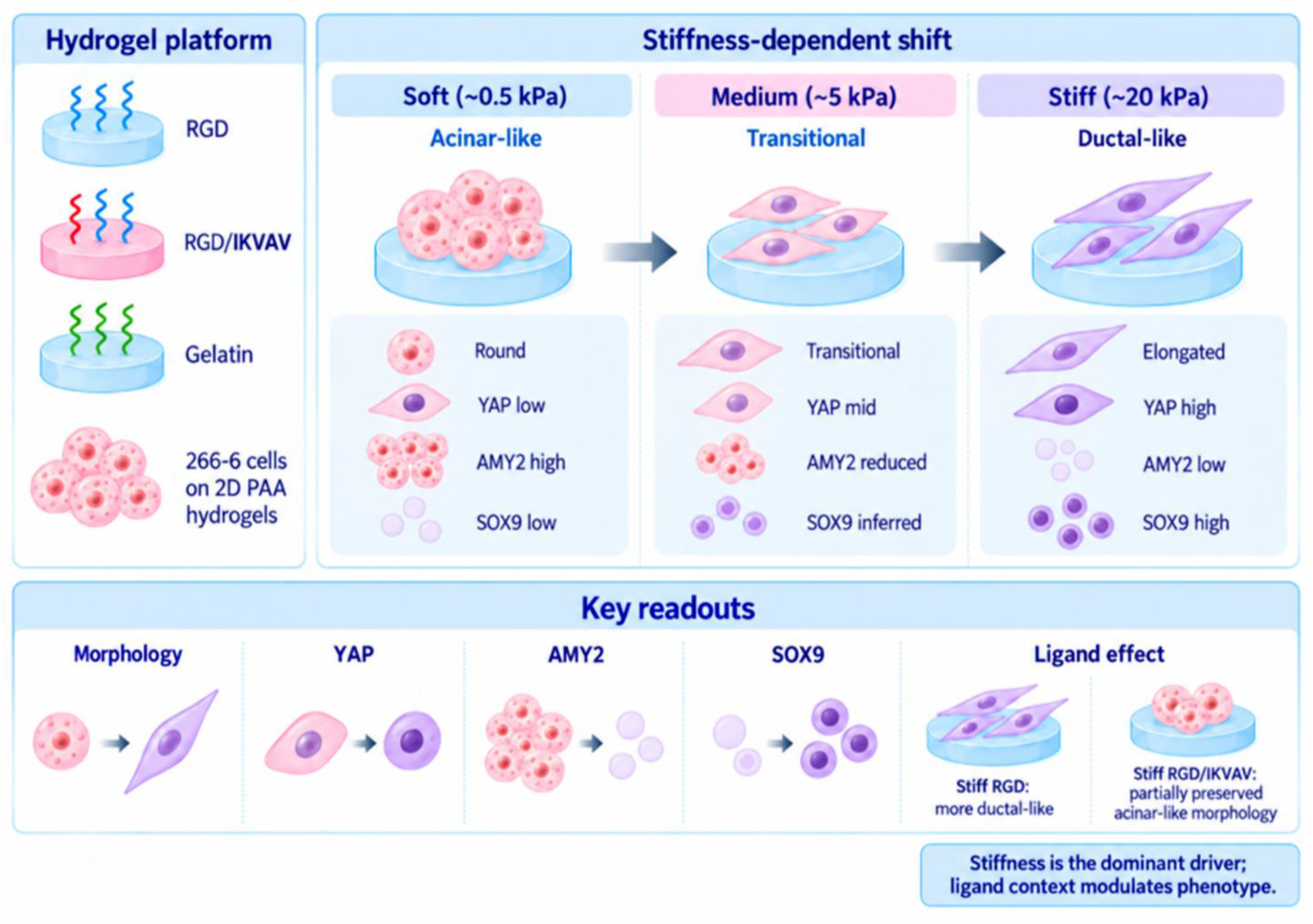
Integrated schematic of matrix stiffness-dependent acinar-to-ductal metaplasia-associated phenotypic changes in the tunable pancreatic hydrogel model. The PAA hydrogel platform enables controlled variation of substrate mechanics and extracellular-matrix ligand presentation for culture of murine pancreatic acinar 266-6 cells. Increasing matrix stiffness produces a progressive shift from a compact, acinar-like phenotype on soft substrates toward an elongated, ductal-like phenotype on stiff substrates, with the medium mechanical condition representing a transitional cell state. Soft substrates are associated with greater cell roundness and geometric regularity, lower aspect ratio, high AMY2 expression, low SOX9 expression, and low nuclear localization of YAP. Increasing stiffness promotes cellular spreading and elongation, increased nuclear YAP localization, loss of the acinar marker AMY2, and acquisition of the ductal marker SOX9. SOX9 was directly quantified in the soft and stiff gelatin-presenting hydrogel conditions; the medium SOX9 state shown schematically represents the overall transitional phenotype inferred from the integrated stiffness series rather than a directly measured medium SOX9 condition. Ligand environments examined in the study included RGD, a mixed RGD/IKVAV condition, and gelatin. Across these conditions, matrix stiffness was the dominant determinant of the ADM-associated phenotype, supporting a central role for mechanobiology in regulating early pancreatic cell-state plasticity. Extracellular-matrix composition nevertheless modulated the phenotypic response: stiff RGD/IKVAV substrates partially preserved acinar-like phenotype despite elevated nuclear YAP localization, indicating that laminin-derived IKVAV cues can buffer aspects of the stiffness-driven phenotypic transition. No direct SOX9 comparison was performed between stiff RGD/IKVAV and stiff RGD or gelatin conditions; the IKVAV buffering effect summarized here is therefore based on morphology together with the YAP and AMY2 readouts. The schematic summarizes the principal conclusion that matrix stiffness drives ADM-associated cellular transformation, while ligand context modulates how this mechanically induced phenotype is expressed.

The schematic was generated using ChatGPT (OpenAI) based on an author-defined conceptual summary of the study findings and was subsequently reviewed and verified by the authors for scientific accuracy.

## DISCUSSION

Several protocol steps are critical for obtaining interpretable results. First, the PAA stiffness must be verified experimentally rather than inferred solely from acrylamide/bis-acrylamide formulation, because small differences in preparation, polymerization, hydration, and batch handling can shift the measured modulus. The main three-stiffness peptide series spans approximately 0.5, 4.5, and 18 kPa, while the adaptable gelatin-functionalized SOX9 implementation uses the soft and stiff mechanical endpoints. This illustrates that the relevant experimental variable is the verified mechanical regime rather than an exact nominal formulation. Second, ligand presentation must be considered independently of stiffness. The FITC-RGD calibration showed that equal soluble peptide inputs yielded higher coupled fluorescence on stiffer PAA, so an uncorrected experiment could confound mechanics with adhesive ligand density. Although the same Sulfo-SANPAH activation and coupling chemistry is used throughout, the gelatin implementation reinforces the need to characterize ligand presentation for each ligand independently rather than assuming that coupled ligand density or coupling efficiency is equivalent across RGD, RGD/IKVAV, and gelatin conditions. Third, passage history, seeding density, and imaging/segmentation rules should be held constant across conditions. The YAP N:C calculation is particularly sensitive to accurate nuclear and whole-cell masks, and the AMY2/SOX9 analysis similarly requires consistent cell-area segmentation because fluorescence is normalized to cell area.

Common problems can usually be traced to polymerization, surface chemistry, or cell attachment. Failure of the gel to remain attached to the coverslip suggests inadequate coverslip silanization, whereas adhesion of the gel to the casting slide suggests insufficient slide passivation. Incomplete polymerization can arise from oxygen exposure or delay after addition of ammonium persulfate and TEMED. If cells attach poorly, first confirm ligand coupling and hydration rather than immediately increasing seeding density. The 50-500 µM RGD optimization performed here showed that increasing ligand concentration improved the proportion of single, spread cells on soft and medium gels, with 500 µM providing the best overall compromise for imaging. The stiffness-specific peptide correction can be recalculated for other PAA formulations, peptide sequences, or coupling chemistries rather than treated as a universal constant. Likewise, alternative ligand environments such as gelatin should be independently optimized for coupled ligand density and cell attachment before direct quantitative comparison with defined peptide-functionalized conditions.

The method has several limitations. The 266-6 line is an immortalized murine acinar model rather than a primary human or primary murine acinar system, and it does not reproduce the full genetic context of PanIN or PDAC, including oncogenic KRAS-driven progression [5]. The two-dimensional geometry also imposes a basal substrate interface and does not reproduce the three-dimensional architecture, fluid flow, stromal interactions, or heterogeneous matrix organization of the pancreas. The peptide-density correction reduced but did not fully eliminate stiffness-dependent differences in FITC-RGD fluorescence, so residual ligand-presentation effects cannot be excluded. Surface topology of the functionalised hydrogels was not assessed, and topographical differences across stiffness conditions could represent an additional uncontrolled variable influencing cell behaviour. SOX9 was quantified as a complementary readout using an adaptable gelatin-functionalized implementation of the same PAA platform using the soft and stiff mechanical endpoints rather than the full three-stiffness RGD/RGD-IKVAV factorial design. This configuration therefore extends the mechanobiological interpretation of the platform but does not establish quantitative equivalence between gelatin and the defined peptide ligand conditions, nor does it directly define SOX9 behavior at the medium stiffness. In addition, the representative study is centered on a day-3 endpoint. AMY2 protein can persist after transcriptional changes, YAP nuclear localization reports mechanotransduction but does not by itself establish irreversible ductal conversion, and a single SOX9 endpoint does not define the temporal order of lineage switching.

These findings are particularly relevant when considered against the broader mechanical evolution of pancreatic disease. Direct measurements of human tissue show that chronic pancreatitis exhibits a higher steady-state modulus than histologically normal pancreas, while PDAC tumours are stiffer still and display distinct viscoelastic behaviour [12]. Although chronic pancreatitis is a recognised risk factor for the development of precancerous pancreatic lesions, only a minority of patients subsequently develop PDAC. More broadly, these observations point to fibrosis-associated mechanical remodelling as a potentially important feature of pancreatic disease. Such remodelling can arise following pancreatic injury in a range of pathological contexts and is not unique to pancreatitis. Rather than representing a defined precursor state, these fibrosis-driven changes may contribute to a tissue environment in which altered extracellular matrix composition and mechanics influence epithelial cell behaviour and potentially modulate the initiation or progression of neoplastic lesions. Reviews from our group have highlighted both the utility of hydrogel systems for reproducing pancreatic mechanical cues and the need to move beyond purely static stiffness toward viscoelastic and mechanically dynamic matrices in cancer and fibrosis [13,14]. Consistent with this, our recent PanIN study used PAA hydrogels to decouple elastic and viscous mechanics and identified viscosity as a driver of EMT-associated phenotypic changes [15]. The present protocol extends this mechanobiological framework to an earlier point in pancreatic disease progression: rather than modeling EMT within an established precursor lesion, it tests whether changes in substrate stiffness and ECM presentation can drive features of ADM, an acinar plasticity event that can precede PanIN formation.

Despite these limitations, the protocol offers a useful advantage over models in which fibrosis, inflammation, ECM composition, and oncogenic signaling change simultaneously. PAA substrates allow elastic modulus to be tuned independently of the adhesive environment, providing a controlled way to ask whether mechanics alone is sufficient to alter acinar phenotype. This reductionist design complements organoid, explant, primary-cell, and in vivo models rather than replacing them. Across the study, increasing stiffness increased nuclear YAP and reduced AMY2 in the peptide-coupled conditions, while adaptation of the same platform to gelatin functionalization and SOX9 quantification showed the same directional phenotypic progression through reduced roundness and geometric regularity, increased aspect ratio, and increased expression of the ductal-lineage marker SOX9 [21,22]. The convergence of YAP activation, AMY2 loss, altered morphology, and SOX9 elevation supports matrix stiffness as a dominant driver of the observed ADM-associated phenotype within the conditions tested in this defined system. At the same time, the retention of acinar-like phenotype on stiff RGD/IKVAV gels despite elevated nuclear YAP illustrates that ECM ligand composition can buffer or reshape how a mechanical signal is phenotypically expressed without overriding the underlying stiffness response.

Future applications can extend the platform in several directions. A day 1–7 time course measuring AMY2, YAP, and SOX9 within the same experimental batches would help resolve the sequence of mechanotransduction, acinar-identity loss, and ductal-lineage acquisition. Direct YAP perturbation by knockdown, constitutively active YAP, or pharmacological inhibition could test whether YAP is required for the stiffness-dependent phenotype rather than merely correlated with it. Because the present study focuses primarily on elastic stiffness, future versions could also introduce independently tunable viscosity, stress-relaxation, or other time-dependent matrix properties to test whether the dynamic mechanics highlighted in cancer, fibrosis, and PanIN [14,15] modulate ADM in addition to Young’s modulus. Additional defined ECM motifs, protein ligands such as gelatin, cytoskeletal-tension inhibitors, mechanosensitive ion-channel perturbations, or controlled inflammatory stimuli could be layered onto the same stiffness matrix. The platform can also serve as a defined two-dimensional screening stage before moving selected conditions into primary-cell, three-dimensional, co-culture, microfluidic, or single-cell transcriptomic models. In this way, the method provides a reproducible framework for dissecting how fibrotic mechanics and matrix composition interact during early pancreatic cell-state plasticity and for prioritizing mechanisms for testing in more physiologically complex systems.

## ACKNOWLEDGMENTS

We acknowledge funding from the StemNiche grant, supported by UK Research and Innovation (UKRI) through the Engineering and Physical Sciences Research Council (EPSRC; grant EP/X036049/1). We also thank colleagues at the Centre for the Cellular Microenvironment (CeMi), University of Glasgow, for technical support provided throughout the study. ChatGPT (OpenAI) was used to assist with the design and generation of the experimental workflow schematic presented in Figure 1 and the schematic presented in Figure 8. The scientific content, structure, and interpretation of both figures were defined by the authors, who subsequently reviewed and verified the final figures for scientific accuracy.

